# Remote Digital Psychiatry: MindLogger for Mobile Mental Health Assessment and Therapy

**DOI:** 10.1101/2020.11.16.385880

**Authors:** Arno Klein, Jon Clucas, Anirudh Krishnakumar, Satrajit S. Ghosh, Wil Van Auken, Benjamin Thonet, Ihor Sabram, Nino Acuna, Anisha Keshavan, Henry Rossiter, Yao Xiao, Sergey Semenuta, Alessandra Badioli, Kseniia Konishcheva, Sanu Ann Abraham, Lindsay M. Alexander, Kathleen R. Merikangas, Joel Swendsen, Ariel B. Lindner, Michael P. Milham

## Abstract

**Background:** Universal access to assessment and treatment of mental health and learning disorders remains a significant and unmet need. There is a vast number of people without access to care because of economic, geographic, and cultural barriers as well as limited availability of clinical experts who could help advance our understanding of mental health.

**Objective:** To create an open, configurable software platform to build clinical measures, mobile assessments, tasks, and interventions without programming expertise. Specifically, our primary requirements include: an administrator interface for creating and scheduling recurring and customized questionnaires where end users receive and respond to scheduled notifications via an iOS or Android app on a mobile device. Such a platform would help relieve overwhelmed health systems, and empower remote and disadvantaged subgroups in need of accurate and effective information, assessment, and care. This platform has potential to advance scientific research by supporting the collection of data with instruments tailored to specific scientific questions from large, distributed, and diverse populations.

**Methods:** We conducted a search for tools that satisfy the above requirements. We designed and developed a new software platform called “MindLogger” that exceeds the above requirements. To demonstrate the tool’s configurability, we built multiple “applets” (collections of activities) within the MindLogger mobile application and deployed several, including a comprehensive set of assessments underway in a large-scale, longitudinal, mental health study.

**Results:** Of the hundreds of products we researched, we found 10 that met our primary requirements above with 4 that support end-to-end encryption, 2 that enable restricted access to individual users’ data, 1 that provides open source software, and none that satisfy all three. We compared features related to information presentation and data capture capabilities, privacy and security, and access to the product, code, and data. We successfully built MindLogger mobile and web applications, as well as web browser-based tools for building and editing new applets and for administering them to end users. MindLogger has end-to-end encryption, enables restricted access, is open source, and supports a variety of data collection features. One applet is currently collecting data from children and adolescents in our mental health study, and other applets are in different stages of testing and deployment for use in clinical and research settings.

**Conclusions:** We have demonstrated the flexibility and applicability of the MindLogger platform through its deployment in a large-scale, longitudinal, mobile mental health study, and by building a variety of other mental health-related applets. With this release, we encourage a broad range of users to apply the MindLogger platform to create and test applets to advance health care and scientific research. We hope that increasing availability of applets designed to assess and administer interventions will facilitate access to health care in the general population.

## Introduction

### Global burden of mental illness and barriers to care

The global burden of mental illness is staggering. Epidemiologic studies indicate that 75% of all diagnosable psychiatric disorders begin prior to age 24 [1] and the lifetime prevalence of a severe disorder among children and adolescents is 21.4%. The most common diagnoses during childhood are anxiety disorders, attention deficit hyperactivity disorder, and mood disorders [2]. Despite their high prevalence, only about 50% of children with a mental health disorder receive treatment (CDC, NIMH). Adults as well as children remain untreated even though effective treatments exist. The World Health Organization has reported large median treatment gaps for alcohol abuse and dependence (78%), generalized anxiety disorder (58%), obsessive–compulsive disorder (57%), depression (56%), dysthymia (56%), panic disorder (56%), bipolar disorder (50%), and schizophrenia (32%) [3,4]. There is a general dearth of mental health resources, especially in low-to middle-income settings [5] and for disadvantaged youth [6], and there are many barriers to care that do not involve finances, insurance, and availability of treatments [6-16]. Simply scheduling an appointment in the U.S. can be difficult [8], and wait times may take months [8], [11]. Additional barriers include concerns about stigma and privacy, as well as exceptional circumstances when telehealth is the only option, such as with the current COVID-19 crisis.

### Increasing need for digital, mobile, remote mental health care and research

People seeking mental health care can use internet-based resources to overcome barriers to treatment such as distance, cost, long waits, inconvenience, stigma, privacy, and problems associated with in-person visits. In recent years, people are increasingly turning to the internet as a preferred source of knowledge about mental health [7,17,18]. However, the abundance of inaccurate and misleading information on the internet makes it difficult to evaluate credibility of information sources, veracity of claims, and effectiveness of recommendations and therapies. In addition, there are many scenarios where obtaining data or care without access to the internet or a computer is important. Furthermore, receiving scheduled notifications at any time and any place (“experience sampling”) can be critical for realistic assessment and effective intervention. In the context of these requirements, mobile devices with dedicated applications offer a better way of receiving vetted, curated content, and of communicating relevant information without having to sit in front of a computer or find the right person to talk to at the right time.

Clinicians and health organizations, particularly those in the mental health sector, are overwhelmed. General health practitioners in particular struggle to serve the needs of their local communities. Better assistance is needed to identify mental health issues during checkups to ensure that these concerns are caught early on and that patients benefit from referral to a specialist. Health organizations struggle to provide broader assistance to the vast majority of people around the world who have limited or no access to mental health resources. They need better tools to scale up mental health efforts that reduce or remove face-to-face interaction to inform, assess, and provide therapy. Digital and mobile tools are especially important to reach people who need care and are in remote areas, socioeconomically disadvantaged conditions, and/or in situations where there is profound stigma surrounding mental illness. Telehealth options that involve video teleconferencing with a therapist are becoming more prominent, as evidenced by new companies that promote internet-mediated talk therapy. However, these services require the time and attention of expensive and limited human resources, and therefore do not scale as well as digital, mobile, and remote resources that do not require human interaction. The current COVID-19 crisis has made it starkly apparent that telehealth, with or without human interaction, may at times be the only option for seeking and receiving care. This pandemic is having a widespread impact on mental health, from the anxiety and stress related to the helplessness, fear, and uncertainties of the crisis itself to the isolation and loneliness of home confinement and social distancing. The need for remotely-administered information, assessment, and therapy will only grow as nations have been forced to discover the extensive possibilities of these novel technologies.

Scientists often struggle to acquire relevant mental health and behavioral data when using traditional research paradigms. Study participant recruitment efforts are usually restricted to individuals who can be evaluated in person and provide written consent, which drastically reduces the size and scope of study samples. For data to be relevant, robust, and replicable, it is important to frequently sample from large and diverse populations, a task which is difficult to accomplish by conventional methods. Off-the-shelf, internet-based tools such as mobile applications for collecting data do not meet the needs of most scientists who must configure their data collection tools to reach the right populations and address specific questions of scientific and clinical relevance. Even tools that are configurable often require considerable mobile application software engineering expertise that is outside of most laboratories’ capabilities. For laboratories that attempt to create their own mobile application from scratch, it is the authors’ experience that they often turn to outsourcing for development of the application only to find that it is far more costly than anticipated, and requires consistent maintenance and upgrading to keep up with changing software dependencies, operating system versions, and hardware, let alone the changing requirements of the research itself [personal communications].

Appropriately-constructed, configured, and vetted mobile applications are currently the most promising avenue for satisfying all of the above needs, as mobile devices with internet access are rapidly becoming widely available to diverse populations around the world. They provide a convenient way to passively or actively collect and present information that can be relevant to the natural variations and social and environmental contexts of people’s lives. Passive methods usually involve sensors carried or worn on some part of the body and can collect objective, real-world data about participants in order to monitor motor activity, sleep, heart rate, cognition, behaviors, mood, and physiological states, as well as detect important outcomes such as medication response, etc. (e.g., [19]).

One important limitation in the interpretation of data acquired through passive monitoring is the lack of contextual information on variables that may influence the sleep, activity, or mood changes inferred from speech, texting, GPS location or other interactions with mobile devices. Although many studies do include subjective symptom ratings with passive monitoring, most of this work is based on retrospective reporting of symptoms over the past week or month. Gaining insight into the directional associations between events and psychological states and their association with sleep and physical activity can be enhanced through concomitant administration of tools that simultaneously capture descriptions of symptoms of mood, cognition, and other subjective experiences, which remain the core components of psychiatric disorder criteria. These active methods of experience sampling include explicit self-reports that may range from occasional and detailed survey instruments to more frequent, brief and in-the-moment questionnaires that are referred to as “ecological momentary assessment” (EMA).

As described by Trull and Ebner-Priemer [20], EMA offers a number of major benefits over traditional clinical assessments including the reduction of retrospective bias, real-time tracking of dynamic processes, simultaneous integration of multi-level data (e.g. biological, psychological), characterization of context-specific relationships of behaviors and symptoms, inclusion of interactive feedback, and enhanced generalizability of results. EMA has been shown to be highly feasible and valid in the assessment of diverse categories of mental illness including patients with mood disorders, anxiety disorders, substance use disorders, and psychosis [21-25] as well as in the assessment of transdiagnostic mental health issues such as suicidal ideation [26]. More recently, EMA has been used to assess cognitive functions (for review, see [27]).

Batteries of cognitive tasks, such as the NIH Toolbox [28,29], ACE [30], Cambridge Cognition [31], and others, are primarily used for research and are not currently intended for clinical practice; they are for the most part commercial, proprietary, and permit limited (if any) configuration options for presenting and collecting data.

Some examples of recent application of sensor-based technologies such as mobile phones and wearable devices have been used to track mental health, most notably through the launch of phone apps such as Northwestern University’s Center for Behavioral Intervention Technologies’ Purple [32,33] and Intellicare [34], [35] platforms, Harvard University’s Beiwe platform [36,37], and apps such as mPower [38,39] built on top of Apple’s ResearchKit for iOS [40], ResearchStack for Android [41], and others. The Beth Israel Deaconess Medical Center maintains an extensive database of apps that claim to respond to mental health needs [42], and provides a protocol for their evaluation [43,44]. There is an unmet need for a free, open, configurable platform to translate pencil-and-paper assessment instruments, cognitive tasks, and therapies into attractive, engaging, and effective digital, mobile, remote tools that exceed current standards of privacy, security, and accessibility.

Here, we (1) review existing digital, mobile, remote tools for configurable data collection and content delivery, (2) summarize the motivation for and development of a new mobile platform called MindLogger, and (3) describe MindLogger applets, including an initial use case as part of a large-scale mental health study.

## Methods

### Review criteria for existing customizable, mobile, experience sampling tools

In the present review, we did not consider apps limited to specific assessments, cognitive tasks, or therapies, but rather examined platforms that enable the creation and distribution of such apps. We wanted to find products that have an administrator interface for creating and scheduling recurring, customized questionnaires, where users receive and respond to scheduled notifications on a mobile device.

For a detailed outline of the protocol we followed, please see Appendix 1, which is briefly summarized here. We gathered information about products with desired characteristics over the last three years through clinical and research collaborators and colleagues. To extend this search we conducted three queries in Google’s search engine (without quotation marks or Boolean operators): (1) “digital electronic data capture systems”, to broadly identify any electronic tools for capturing data; (2) “mobile phone software sensor data collection”, to identify mobile data collection software that may involve sensors; (3) “alternative to qualtrics”, to identify alternatives to one of the most prominent products that enables online customization of surveys (though Qualtrics itself does not currently have a mobile app). We visited the websites of the first 20 search results from each query (not including advertisements) and identified candidate products. For example, if a website listed the “Top Ten Apps for…”, we would include these ten apps in our initial set of candidates. We then filtered this set by the following inclusion criteria: the candidate product had to (a) be in current use, (b) have (Android and iOS) mobile apps, and (c) have an administrative user interface for creating and scheduling times and days for recurring, customized questionnaires, where end users receive and respond to scheduled notifications via an iOS or Android app on a mobile device. Where there was any ambiguity, we contacted the company/organization to clarify, scheduled an online demonstration, and requested a free trial to explore the product. We excluded products that do not currently fulfill the above requirements or for which we could not receive a demonstration or trial without a legal agreement. We also excluded products that require SMS, email, or other modes of communication outside of their mobile application to send and receive notifications. Since the type of notification is important in experience sampling applications, we identified which products can deliver local operating system notifications or push notifications, where an end user receives notifications in their mobile device’s notification bar at scheduled times, and a tap on a notification takes them to their scheduled activity within the mobile app. Local operating system notifications do not require an internet connection at the time that the notification is to be received, whereas push notifications do, and both of these are distinct from simple in-app notifications, which require the end user to be using the app to see their notifications.

We collected additional information from product websites, teleconferences and online demonstrations with the product creators, and from free trials, to determine the degree to which each is (a) customizable in its information presentation and data capture capabilities, (b) private and secure, and (c) accessible (easy to use, economical, and open source). Customization is important to ensure that content, language, and presentation can be adapted and updated to meet the specific needs of a given population of intended end users, and to expand the scope of possible dimensions to assess, analyses to perform, and inferences to make. Privacy and security is a central concern, especially as we conduct research involving a triply vulnerable population of end users: (i) child and adolescent (ii) patients with (iii) mental health and learning disorders. Data access, encryption, and deletion capabilities are the primary considerations in this domain. “Accessible” can mean many things; here we refer to whether and how the administrator can access the product, its source code, or its data. The degree to which a product is affordable will often determine the degree to which it is adopted.

Possibly the most stringent accessibility criterion one could have is for the software to be open source. Open source software is important because the product does not live or die with a given company, provides an opportunity for anyone to build on and improve the software, and it is open to greater scrutiny to ensure quality of the software, accuracy of any claims made about the software, and transparency of clinical and scientific practices that use the software. Each of the above characteristics offers a competitive advantage, but they should be considered together. For example, a company offering a free product could store or transmit user data insecurely.

### Development of MindLogger

MindLogger [45] is intended as an *open ecosystem to create, edit, share, and administer mobile/web “applets” for data collection and content delivery*. We use the term “applet” to refer to a customized collection of activities within the MindLogger app administered to target end users. Our focus with MindLogger is to easily create and edit digital mental health assessments and interventions and administer them to users via mobile or web applets. The three key innovations we sought to accomplish with MindLogger were: **customizability** of content, response options, and appearance; an extensive library of applets built using **open standards** (open, reusable parts defined by an open protocol); and distribution as a **single app** that appears differently to different user groups.

MindLogger development began in 2017 with a focus on mental health assessments. After significant development of an early prototype to present to mental health research and clinical colleagues at the Child Mind Institute and the Child Mind Medical Practice in New York City, we revisited development with a greater emphasis on human-centered design [46,47]. We included a variety of key stakeholders to ensure that their needs would be met by the platform: clinicians (psychiatrists, psychologists, social workers, etc.), scientists (neuroscientists, cognitive psychologists, etc.), and directors of schools specializing in learning and developmental disorders, as well as technology consultants. We integrated their feedback at different stages throughout the development of the platform.

### Application of MindLogger in the Healthy Brain Network study

The Child Mind Institute’s Healthy Brain Network study [48] is an ongoing initiative focused on creating and sharing a biobank of data from 10,000 New York area participants (ages 5-21). The Biobank houses data about psychiatric, behavioral, cognitive, and lifestyle phenotypes, as well as multimodal brain imaging (resting and naturalistic viewing functional MRI, diffusion MRI, morphometric MRI), electroencephalography, eye tracking, voice and video recording, genetics, and actigraphy. In order to obtain more in-depth information on real-time tracking of emotions, behavior, daily activities, and their contextual influences, we have adapted the the combined actigraphy and EMA mobile assessment tools and content from the National Institute of Mental Health (NIMH) Family Study of Affective Spectrum Disorders, a large community-based controlled family study [49]. The original EMA collected data four times per day for two weeks from phones provided to study participants (the Tungsten E2 model of a Palm mobile device and more recently from Android mobile phones). See below for an excerpt from a description of its content:

> “The EMA assessments included questions concerning a diversity of daily life experiences and behaviors, including data assessed at the moment of the EMA signal (current location, social company, performance of specific behaviors, mood states) and data assessed over the time period between the current and previous assessment or, for the first assessment of the morning, since awakening (experience of daily events and event negativity, food intake, substance use, experience of headache and its specific symptoms). Additional questions were asked at the first EMA assessment of the day concerning sleep duration, quality and sleep problems, and at the end of the day concerning global ratings of the stressfulness of the day, food craving for the day and specific physical symptoms (gastrointestinal symptoms, muscle pain). The response possibilities included Likert scales for dimensional constructs (such as mood or event negativity) and checklists that allowed for either multiple responses (such as for noting all food types consumed since the last assessment) or single responses (such as current physical location). …For daily events, participants were asked at each assessment to identify the one event or experience, good or bad, that had affected them the most since the last questionnaire, and to rate the impact the event had on them on a 7-point Likert scale…”

The EMA data from the NIMH Family Study that evaluated the association between daily events and emotional experience yielded important differences in patterns of reactivity among the major subtypes of mood disorders, including bipolar I disorder, bipolar II disorder, major depression, anxiety disorders without a mood disorder, and controls [50]. These findings demonstrated how EMA is a particularly well-adapted tool for assessing affective dynamics as well as emotional reactivity following daily life events. The value of combined passive and active monitoring in this study further showed bidirectional associations between energy, motor activity, and sleep, and unidirectional associations between activity and mood, suggesting that increased activity could be used as an intervention for depression [51]. Using the novel analytic approach of fragmentation for testing the stability and instability of emotional states in this study showed greater instability of energy and attention in people with a history of bipolar I disorder, whereas those with bipolar II disorder or major depression exhibited greater fragmentation of mood and anxiety [52]. Although these findings were primarily based on adult samples, the inclusion of a substantial subset of offspring ages 10-18 of parents with mood disorders and controls provided compelling evidence for the feasibility, acceptability, and clinical significance of EMA in youth. Therefore, the goal of the present initiative was to create a version of the EMA in MindLogger, with updated content (particularly with regard to sleep, positive/negative thoughts, food/drink, internet, and social media questions), enhanced with clarification of the content, inclusion of colorful images, and formats adapted for children and young adults to encourage engagement [53]. MindLogger’s “NIMH-EMA” applet has been launched as part of the Healthy Brain Network study and is now being built into data collection of the NIMH research program.

## Results

### Comparison of reviewed experience sampling tools

Our search resulted in 392 products, of which 315 appear to be in current use. Appendix 1 contains a list of 101 of these products that have Android and/or iOS mobile apps. Upon closer inspection of their websites, 59 appear relevant to scheduling questionnaires and notifications for a group of respondents, so we contacted the 59 products’ companies/organizations through their online contact forms or via email to clarify their products’ capabilities, and exchanged emails with the 47 companies that responded. Based on these exchanges, we were able to identify 21 products that appeared to satisfy our primary criteria (administrator interface for creating and scheduling recurring, customized questionnaires, where users receive and respond to scheduled notifications on a mobile device). Five potential candidates were omitted because they are intended for use by internal business employees of a company, not by participants of a study, and require individual licenses or logins or fees per device. Two more candidates were omitted because they required a legal agreement to demonstrate their product. Of the 14 remaining products for which we engaged in demonstrations and free trials, 10 currently meet primary criteria and are included in Figures 1-3.

**Figure 1.**
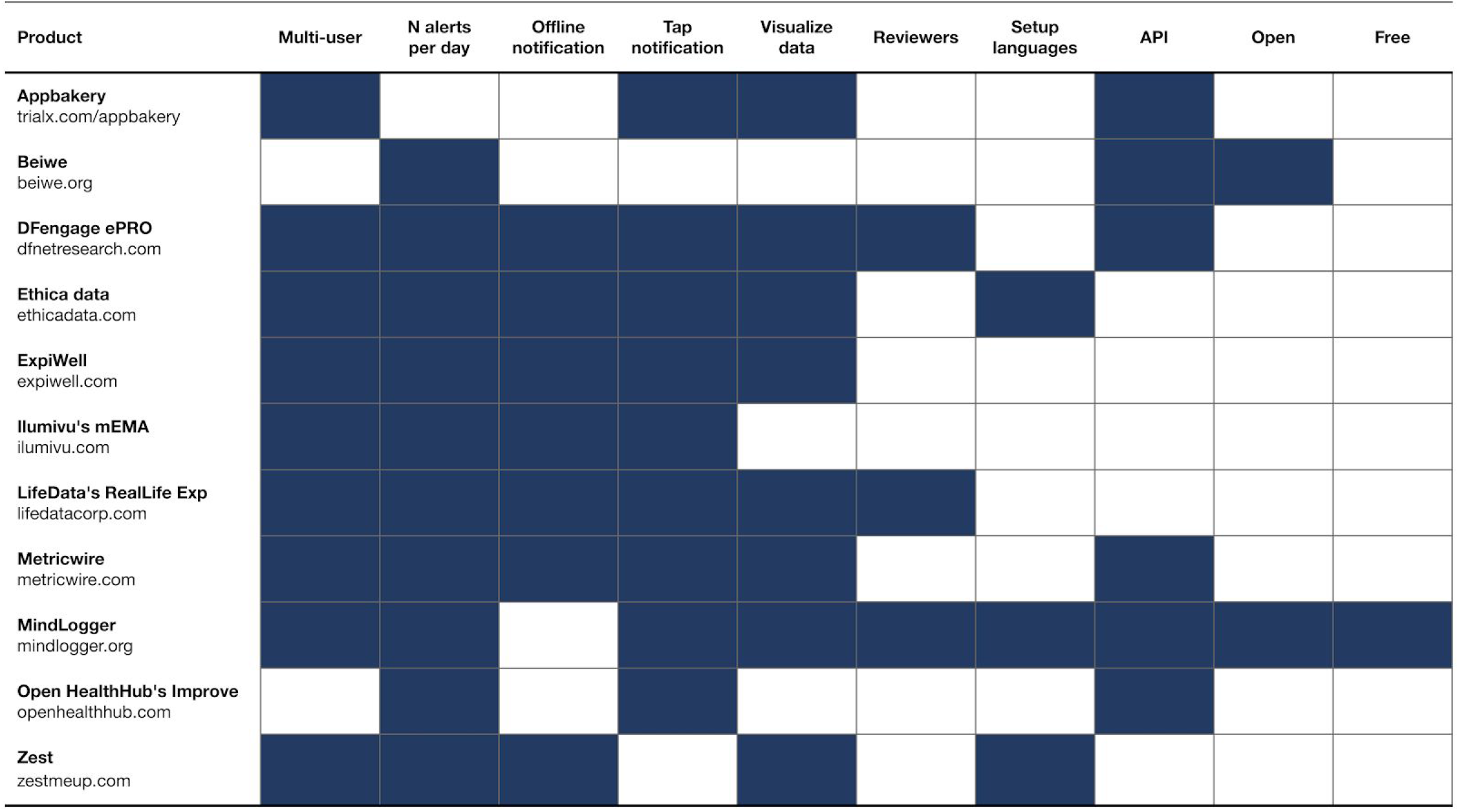
Access to experience sampling products, software, and data.

Figure 1 contains information about access to the product, software, and data (for both Android and iOS):

- **Multi-user:** Can more than one end user access the app on the same mobile device, even if it means logging out and logging back in again?
- **N alerts per day:** Can end users receive more than one notification per day per activity?
- **Offline notification:** Can end users receive and respond to any notifications without an internet connection?
- **Tap notification:** When an end user taps on a notification in their mobile device’s notification bar, does it take them directly to their scheduled activity within the app?
- **Visualize data:** Is there a data visualization dashboard to review any individual end user’s response data?
- **Reviewers:** If there is a data visualization dashboard, can an administrator give someone access to review only one end users’ response data in the dashboard?
- **Setup languages:** When administrators create a customized questionnaire, can they choose from at least five different languages to use the interface, in addition to English? (This is distinct from how many different languages the end users can see.)
- **API:** Is there a consumer-facing application programming interface (API)?
- **Open:** Is the product’s mobile application software fully open source?
- **Free:** Do you have to pay money to use the product in a way that satisfies our primary criteria for personal or research purposes?

Figure 2 contains information about presentation and data capture features:

**Figure 2.**
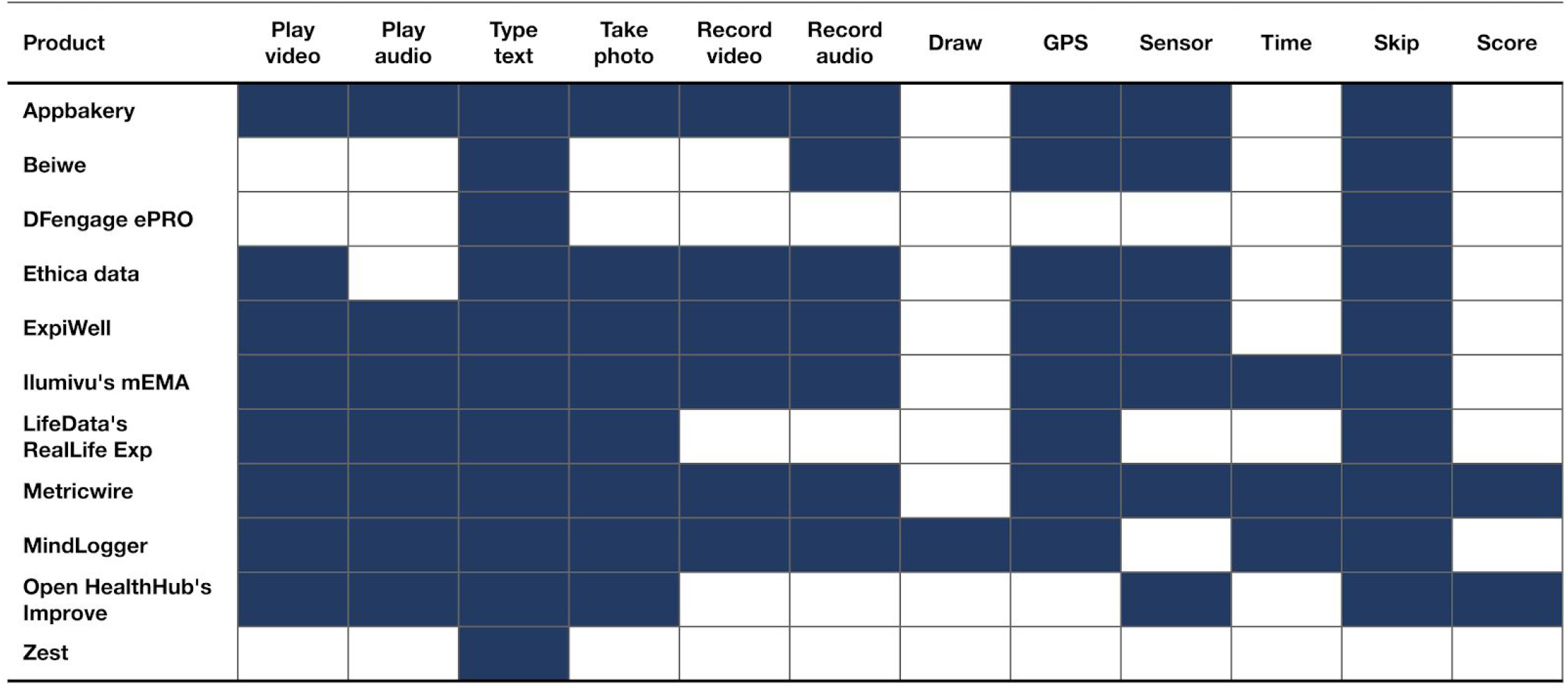
Presentation and data capture features of experience sampling products.

- **Play video/audio:** When administrators create an activity, can they include video or audio clips?
- **Type text, Take photo, Record video/audio, Draw:** When administrators create an activity, can they include any of the following to capture data from end users?: text entry, camera photo, audio/video recording, drawing
- **GPS:** Can GPS location data be acquired through the app?
- **Sensor:** Can any additional sensor (e.g., accelerometer) data be acquired?
- **Time:** Can any question include a countdown/timer?
- **Skip:** Can the response to a question determine which is the follow-up question (skip/branch logic)?
- **Score:** Can a questionnaire’s scoring logic be entered when creating the questionnaire?

Figure 3 contains information about privacy/security:

**Figure 3.**
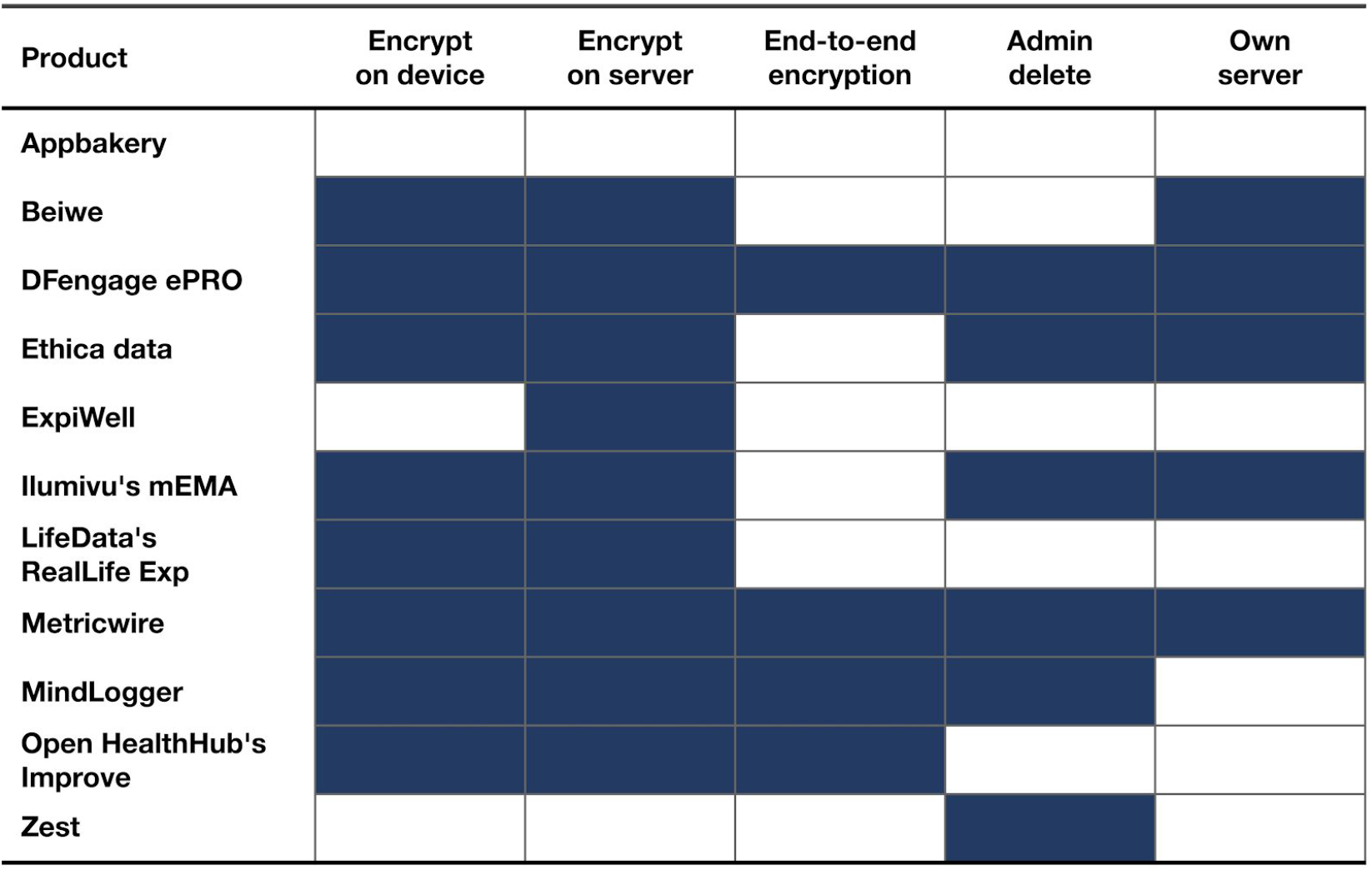
Privacy and security of experience sampling products.

- **Encrypt on [device, server]:** Does the product encrypt data on the mobile device or on the server?
- **End-to-end encryption:** Is the product end-to-end encrypted, or could someone within the company or organization hosting applets on their server see (even anonymized respondents’) response data?
- **Admin delete:** Can an administrator delete an individual end user’s data without having to make a request from the product creator?
- **Own server:** Can the product be hosted on an administrator’s server?

### MindLogger platform

#### Roles and permissions

Figure 4 shows a schematic of MindLogger, where an administrator selects, edits, or creates an applet, administers the applet to end users, and views or makes use of their data. Figure 5 outlines the different roles (owner, manager, coordinator, editor, user, reviewer) and their permissions with regard to administration, content, use, and data management.

**Figure 4.**
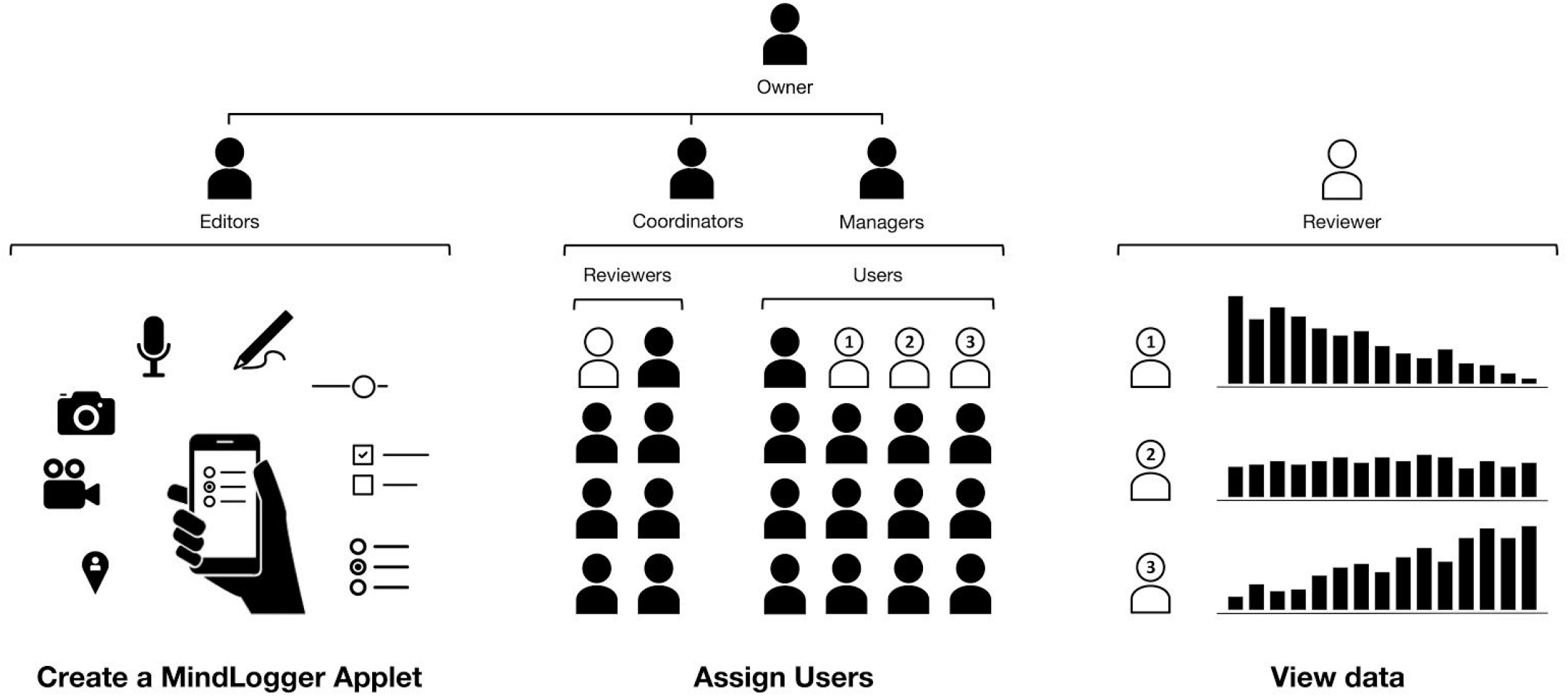
MindLogger schematic. Left: Select, edit, or create activities with scheduled notifications, such as questionnaires, tasks, or interventions, for use on mobile (iOS, Android) devices or the web. Middle: Assign yourself or others to do these activities for data collection, annotation, research, or remote clinical assessment or therapy. Right: View end user data for which you have access.

**Figure 5.**
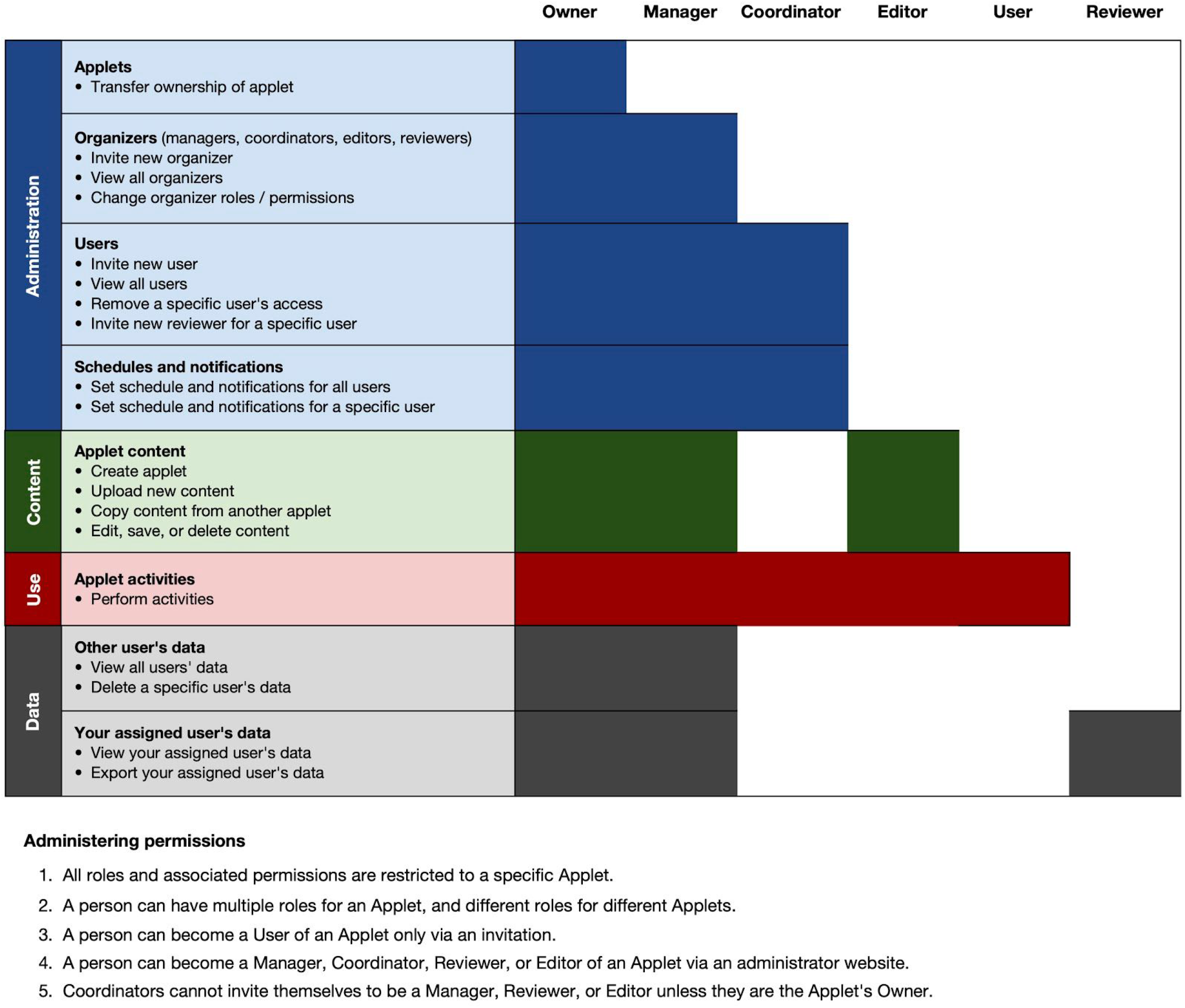
MindLogger roles and permissions for administration, content, use, and data management.

#### MindLogger architecture

MindLogger’s architecture (Figure 6) consists of a set of end-user-facing front-ends (two mobile applications and a web application) and organizer-facing front-ends (an admin panel, data dashboard, and applet builder) with a shared RESTful HTTP API using MongoDB [54] for data storage. The mobile front-ends, Android and iOS apps, are built using React Native [55]. This allows us to share a single code base across mobile platforms, resulting in increased speed of development and ease and cost-effectiveness of maintenance. The web application is a ReactJS browser-based counterpart to the mobile applications, and currently provides a subset of their functionality. Organizers (managers, coordinators, editors, and reviewers) have access to different single-page applications built using VueJS [56]. The admin panel and applet builder are for managing user roles and applets, and the data dashboard is for reviewing user data, with custom charts implemented using d3.js [57]. The computer security firm Alpine Security [58] has conducted extensive cybersecurity black, gray, and white penetration tests to ensure that MindLogger follows best practices for privacy and security.

**Figure 6.**
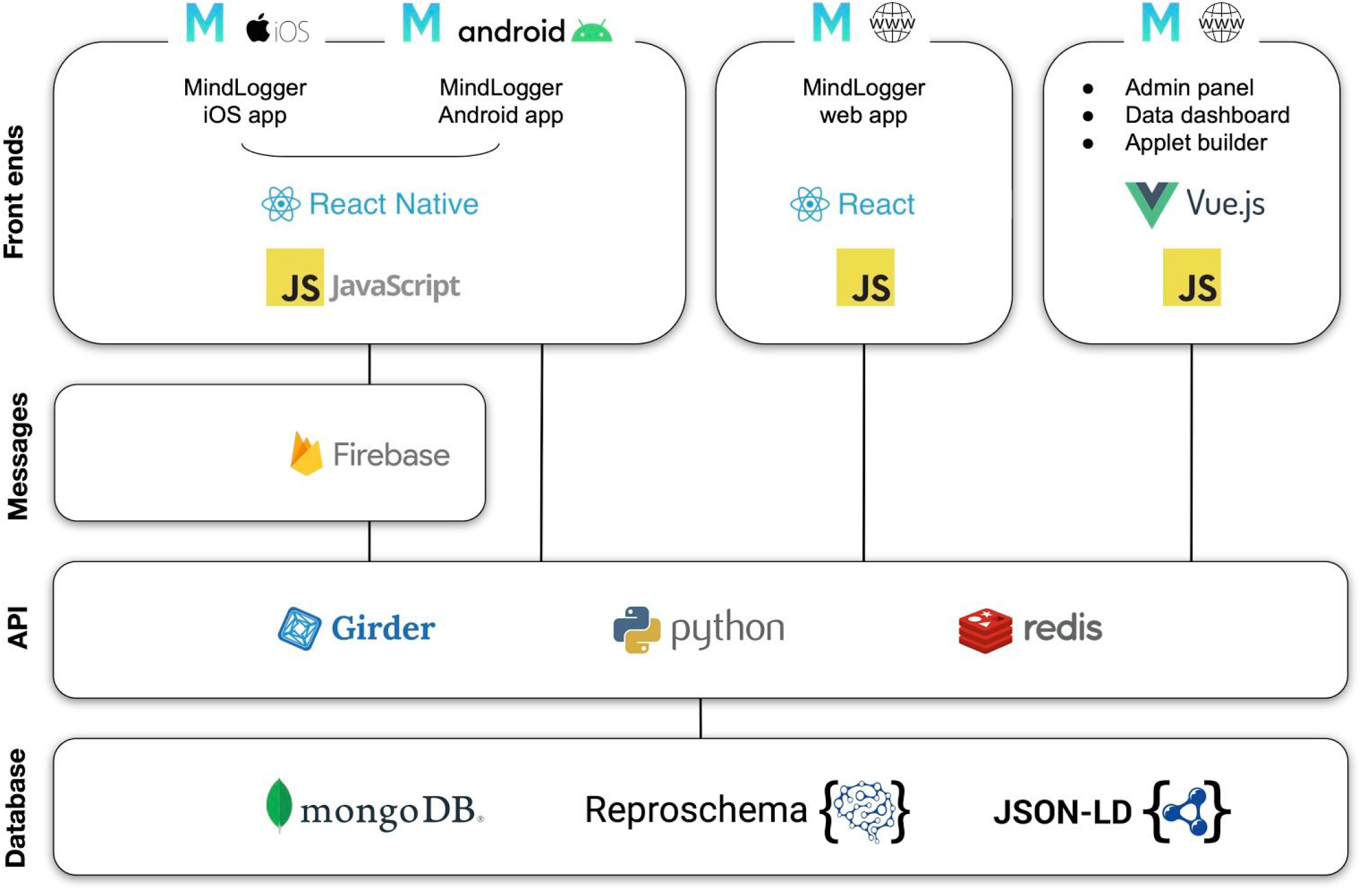
MindLogger architecture diagram.

Our mobile app front-end and back-end code base is accessible as online GitHub repositories [45], and is licensed under an extremely permissive, Open Source Initiative-approved open-source license, the Common Public Attribution License (CPAL-1.0) [59]. The license requires that attribution be given by including (1) the copyright notice: “Copyright (c) 2017 MATTER Lab at the Child Mind Institute”, (2) the URL: https://matter.childmind.org“, (3) the Child Mind Institute’s logo, and (4) the attribution phrase: “Child Mind Institute product intended for building applications for good.” We include the attribution phrase to give credit to the developers while also making it clear that while we intend for people to build MindLogger applets that will be benevolent, we have no control over their intent, content, or the data they collect. See [60] for an example license file.

The back-end API is built in Python using Girder’s RESTful API [61] with the CherryPy framework [62]. This software layer provides a set of RESTful endpoints that allow you to manage users, applets, activities, items (such as individual questions), and user responses. All user data is stored in a MongoDB database hosted in an Amazon Web Services [63] cloud instance with password-based encryption. Specifically, user response data is encrypted using their own password on the client side so that only managers or reviewers can view their data using an applet password; other sensitive information (name and email) is encrypted on the server side. We have HIPAA compliance agreements with Amazon Web Services, Google Cloud Platform [64], and MongoDB Atlas [65], and we will support arbitrary backend servers in the near future. For improved performance, MindLogger uses a Redis [66] instance as a temporary storage for data caching. MindLogger uses Firebase Cloud Messaging for sending notifications from the back-end server to the end user’s mobile device. All additional data is consumed from the back-end API through HTTPS requests.

The applets, activities, and items are described using ReproSchema [67], an emerging standard to capture and harmonize cognitive, clinical, and behavioral assessments and responses in a provenance-preserving manner. The schema uses JSON-LD [68] as its representation format and captures, as a connected graph of information, the details of the questions, presentation logic based on responses or scheduling, computation of scores, and interface hints for applications such as MindLogger. The schema uses GitHub to maintain versions and provide persistent URIs for applets and activities, supports multilingual applets, and uses W3C-PROV [69] to establish provenance between the response, the responder, and the applet.

#### MindLogger current features

We have succeeded in implementing the following features (see the MindLogger website [45] for updated information on the project, including features, installation, administration, and use):

- Any of a wide variety of user response options per screen: buttons, boxes, sliders, audio, camera, drawing, location, etc., with response delay and timer options
- Arbitrary number of screens per activity, with user response-based conditional logic directing the sequence of screens
- Arbitrary number of activities per applet
- iOS, Android, and web browser compatibility
- Browser-based administration panel for user management
- Scheduling of applets and notifications per activity per user or group of users
- Browser-based applet builder to easily customize one’s own mobile/web applications without programming or design experience
- Browser-based visualization dashboard for viewing and exporting data
- Open source code, apps, and applets free for personal use
- End-to-end encryption

Each screen of a MindLogger applet can display text and a picture or video, play a sound file, as well as present an interactive component with different possible response options, such as: single- or multiple-selection check boxes, image selection, slider bar, text entry, table text/number entry, audio recording, photo/video capture, drawing/tapping on images, or GPS location button. Figure 7 shows screenshots of MindLogger mobile phone app features. We have created applets to remotely administer assessments as well as therapies, and are currently constructing a public library from over 100 mental health and cognitive assessments that have open licenses for general use.

**Figure 7.**
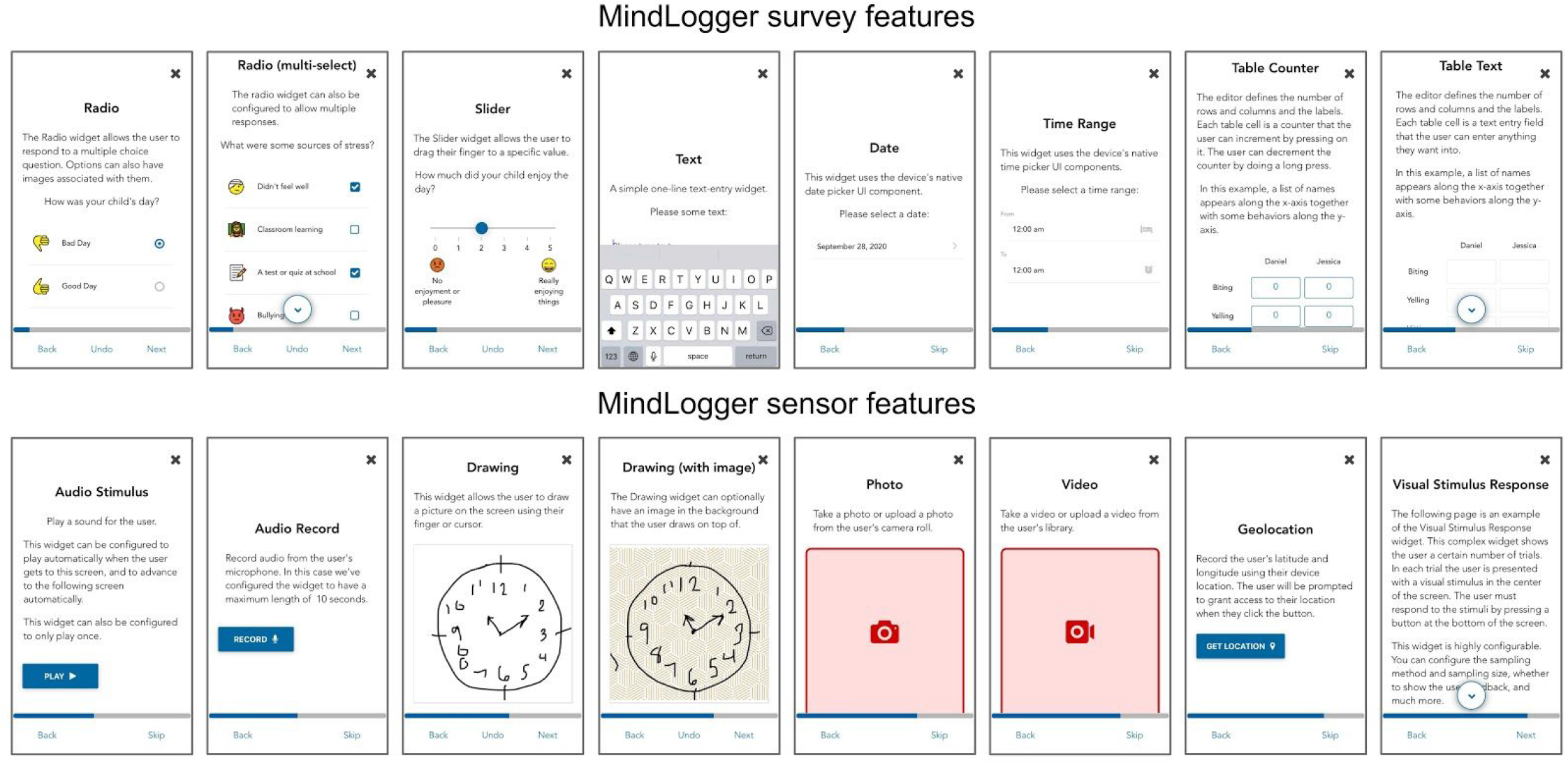
MindLogger screenshots showing survey and sensor features.

### MindLogger NIMH-EMA applet in the Healthy Brain Network study

MindLogger currently administers assessments in our NIMH-EMA applet as part of the Healthy Brain Network study [48] to a vulnerable transdiagnostic New York City community sample (current n=4,315; enrollment rate: 90 per month; >90% have mental health or learning disorders). Study participants receive multiple notifications per day on their Android or iOS device to respond to morning, afternoon, and evening assessments. Figure 8 shows screenshots of the NIMH-EMA applet. We are currently enrolling children and adolescents who are at least 11 years old to use the NIMH-EMA applet as part of the Healthy Brain Network study.

**Figure 8.**
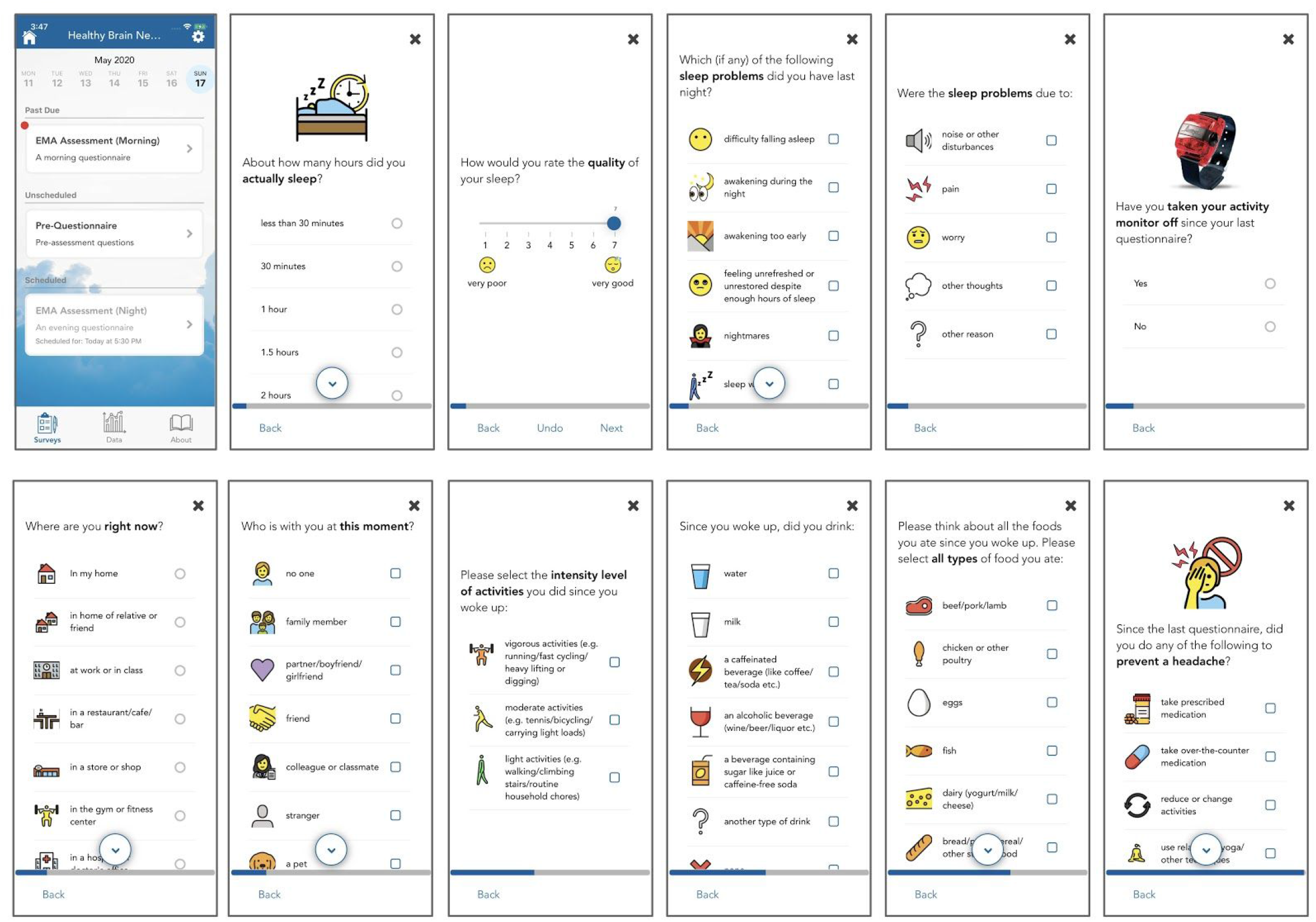
MindLogger NIMH-EMA applet screenshots.

#### EDUCATE study’s daily and weekly diary applet

Having developed and deployed the NIMH-EMA applet as part of one research study, we were able to rapidly develop and deploy a second assessment applet as part of a research study about reading disability. Reading disability is the most common learning disability, affecting 10–15% of school age children [70]. It incurs major functional impairments at all stages of life. A wealth of data documents lifelong disadvantages in educational and occupational attainment. Current evidence-based reading interventions largely rely on services by trained specialists, either in well-resourced classrooms or clinical settings. As such, under-resourced schools (or countries) often are unable to provide reading interventions for their students. The significance of this dilemma is compounded when considering that children of lower socioeconomic status, and children with other serious comorbid behavioral health conditions, may have more severe or complex reading disability profiles [71-73]. Thus, the children most in need are the least likely to have access to evidence-based treatment.

The EDUCATE study, funded by the National Institute of Child Health and Human Development, is a collaborative effort between researchers at the University of Connecticut and the Child Mind Institute. This clinical trial will examine the effectiveness of an at-home, game-based intervention for reading disorders. Parents of participants in this study will complete daily and weekly assessments in MindLogger, which will allow researchers to assess the home environment and compliance with the intervention protocol throughout the study period. This data will be critical for evaluating the impact of this clinical trial.

### MindLogger applets in preparation for deployment

We are currently refining and testing MindLogger applets to assess and administer interventions targeting specific subgroups of youth with particular mental health and learning disorders. While some of these applets support specific collaborators’ research, others are for broader use (listed below, with video screencast demonstrations on the MindLogger website [45]).

#### Pediatric Screener mental health screening applet

Integrating primary care and mental health has been associated with improved patient outcomes [74], so mental health screening in pediatric clinics could lead to earlier diagnosis and improved outcomes for patients. The Hearst Foundations are supporting development of a pediatric screener tool using MindLogger. This tool will administer assessments to children or their parents, for children receiving a wellness checkup at their pediatric clinic, and alert their doctor if a child shows signs of a mental health disorder. We have built a prototype of the applet and will pilot it at the Richmond University Medical Center in Staten Island, New York. The initial screening questionnaire assesses internalizing and externalizing symptoms, issues of attention and hyperactivity, depression and suicidal ideation, disordered eating behavior, and experiences of bullying. It also collects demographic information about the child and parent. Children with a clinically-significant level of symptoms are prompted to complete additional questionnaires to collect more detailed information. We have piloted a similar questionnaire in several New York City-based pediatric settings and have found it to be an effective tool for identifying children at risk of a serious mental health disorder. The MindLogger platform will create a much more streamlined process and user-friendly experience, increasing the probability of adoption by more pediatric practices and clinics.

#### Dialectical Behavior Therapy (DBT) applet

The DBT Diary Card is a digitized version of the diary card used in evidence-based DBT programs. This tool is a daily tracker of mood, targeting behavioral urges and specific behaviors and using coping skills to manage these emotions, urges, and behaviors. Our Dialectical Behavior Therapy applet (Figure 9) will enable therapists to create a digital DBT diary card with a patient to include specific treatment targets for that individual. It will allow patients to set notification reminders to complete the diary card on a daily basis. The patient’s device will automatically update with the new targets and schedule, allowing the user to progress in the therapy without having to change the way the data is retrieved. Both patients and therapists will have access to the diary card data to guide treatment planning and sessions.

**Figure 9.**
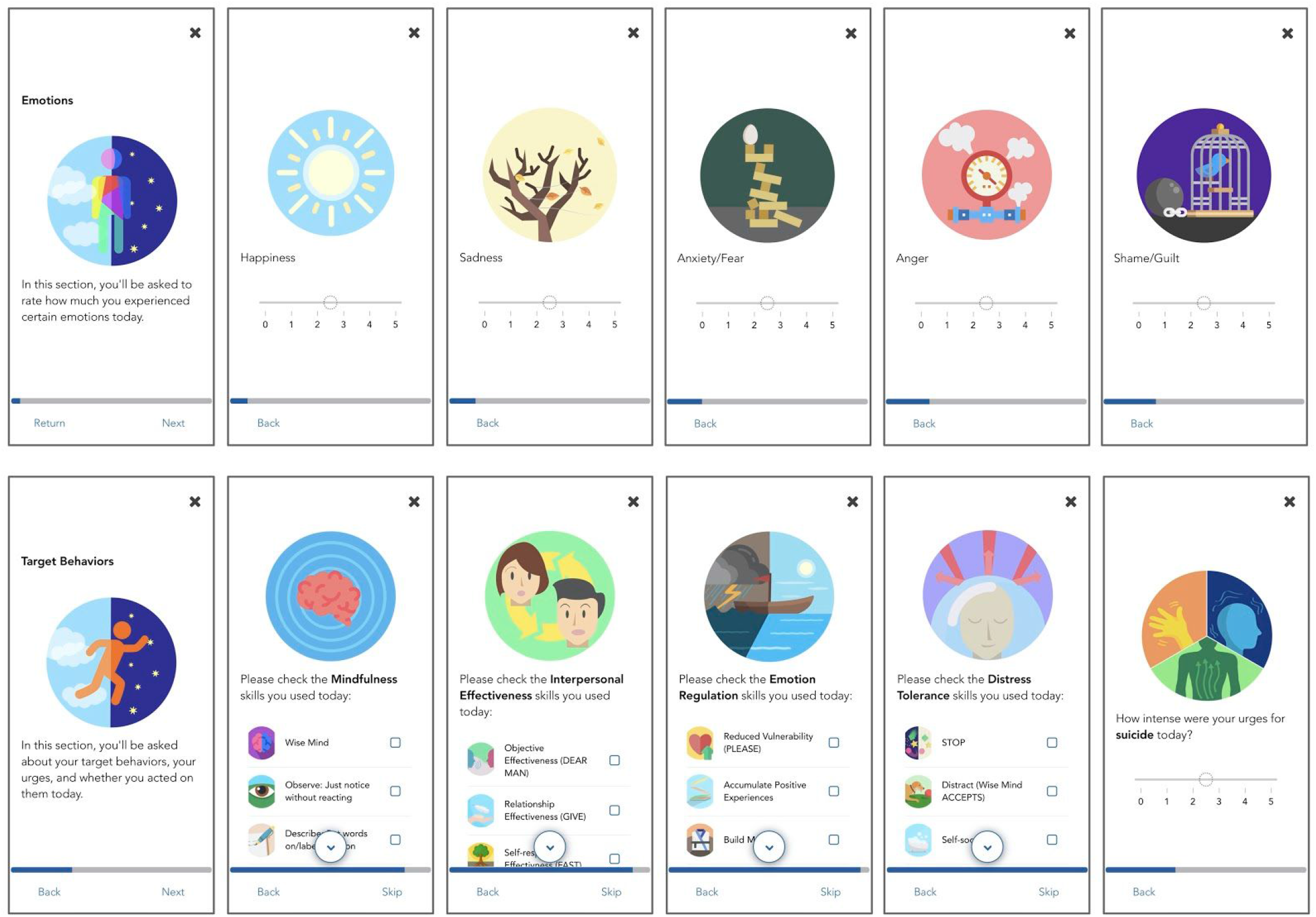
MindLogger Dialectical Behavior Therapy applet screenshots.

#### TokenLogger behavior intervention applet

We are refining a behavior intervention applet called “TokenLogger” that can be customized to track specific behaviors and help parents implement an evidence-based token economy that promotes and reinforces positive behaviors while reducing or extinguishing negative behaviors. It will also help track the frequency, duration and severity of target behaviors to inform modifications in behavioral treatment plans and assess progress towards outcomes and goals. The TokenLogger applet will be evaluated by clinical experts at the Child Mind Institute and piloted with patients at the Child Mind Medical Practice, to better understand the timing, duration, and frequency of undesirable behaviors, and to test the efficacy of this digital rendition of behavior modification therapy.

### Planned MindLogger applets

We intend to replace all of the Child Mind Institute’s and the Child Mind Medical Practice’s pencil-and-paper assessments with MindLogger applets and, where appropriate, create and test digital renditions of therapies as MindLogger applets, and make these universally accessible as part of an online public library of MindLogger applets for anyone to use, modify, and translate. In addition to these, below are a few applets in the planning stages.

#### Diagnostic screening applet for Kiddie Schedule for Affective Disorders and Schizophrenia (KSADS)

We are currently developing a MindLogger applet with part of the Composite International Diagnostic Interview Screener that has been incorporated in the NIMH version of the epidemiologic version of the Kiddie Schedule for Affective Disorders and Schizophrenia (KSADS). This screener is being tested for parent and self-administration in order to streamline the process as we transition to more automated approaches for large-scale studies of youth.

#### Taction exposure therapy applet

We created a prototype iOS and Android mobile app called “Taction” that is a simple exposure therapy game for children who suffer from obsessive-compulsive disorder and/or anxiety-related issues. The app rewards users for tapping on images that heighten their anxiety, potentially helping them progress in their treatment between exposure therapy sessions. Once incorporated into MindLogger, the Taction applet will be evaluated at the Child Mind Institute and piloted with patients at the Child Mind Medical Practice to assess the relative efficacy of a digital rendition of exposure therapy.

#### NIMH cognitive task battery

In the same manner as we want to enable anyone to create, configure, and administer their own mobile questionnaires, we also want to enable anyone to do the same for different types of cognitive assessments. In this spirit we created a Flanker task applet that is thoroughly configurable (presentation and timing of a fixation target, stimulus, and feedback), and we are collaborating with the NIMH to create a small battery of cognitive tasks for research in modeling behaviors.

## Discussion

We reviewed existing mobile data collection platforms, discussed the motivations for and development of the MindLogger platform, and described some of our MindLogger applets, including the initial use case in a large-scale study. Our review is not exhaustive with regard to either the products that could possibly be used for experience sampling or the range of features that these products support. Existing products include an impressive assortment of non-overlapping features and complement each other in ways that reflect the different niches/markets for which they were intended. Limitations of our search for existing tools include: (1) our search queries and Google’s searching algorithm may not reflect the optimal search criteria for some relevant products; (2) product websites are sometimes unclear and incomplete and may even misrepresent the capabilities of their products; (3) some companies did not respond to questions even through their own website’s question/support page. Products are also adding features over time, and some companies or organizations offer paid services to build in desired features. Our comparison should be seen as a snapshot of the current state of a subset of features provided by software products intended for experience sampling.

MindLogger is a new platform that meets most of the stated needs of our collaborators around the world, who desire an open source mobile mental health platform for informing, assessing health, acquiring data, and administering therapies. We prioritized clinicians’ and researchers’ needs and users’ experience during development, aligned technologies to meet these priorities, and are on track to achieve the full feature set we set out to include. One limitation of MindLogger is that it currently does not support passive data collection, or interaction with peripheral devices. This was intentional to reduce concerns about surveillance in an app whose first use-case was for assessing children and adolescents. However, in the future we do intend to support passive monitoring of location and behaviors, and communication with other devices, so long as these are opt-in by the end user and there are clear reminders to the end user concerning what data are collected and how they will be used. Another potential limitation lies in its core strength: By making MindLogger flexible, modular, cross-platform, and configurable to help meet the unforeseen needs of future applet builders and users, the creation of variants of even well-vetted instruments is likely and will necessitate their careful validation.

## Acknowledgments

This project was generously supported by the Hearst Foundations, the Intramural Research Program of the National Institute of Mental Health (grants with PI KRM: ZIA MH002954 Motor Activity Research Consortium for Health (mMarch) and ZIA MH002804-15 Family Study of Affective and Anxiety Spectrum Disorders), the Centre for Research and Interdisciplinarity (CRI) - Université de Paris, the Blanca and Sunil Hirani Foundation, MIT through the laboratory of SSG (SSG and SAA were supported by grant: NIH P41 EB019936), and by Joseph Healey. We thank the companies and organizations that responded to questions about and provided demonstrations of their products. We thank Rangle.io for their help in refactoring MindLogger’s mobile app code base and Alpine Security for conducting penetration tests of the entire code base. We thank Paul Mitrani and Stephanie Samar for clinical feedback during the development of MindLogger applets, and volunteers Chynna Levin, Mairav Linzer, and Benjamin Vogel. Finally, we would like to thank all of our colleagues and collaborators around the world who have encouraged and helped guide the development of MindLogger in anticipation of using it for their many different data collection, research, clinical, and other endeavors!

A Krishnakumar was affiliated with the MATTER Lab at the Child Mind Institute and with the Centre for Research and Interdisciplinarity (CRI) - Université de Paris at the time of this study and is currently affiliated with the ETH Library Lab, ETH Zurich and Citizen Science Centre in Zurich. JC and A Keshavan were affiliated with the MATTER Lab at the Child Mind Institute for part of this study; JC is currently affiliated with the Computational Neuroimaging Laboratory at the Child Mind Institute, and A Keshavan is currently affiliated with Octave Bioscience.

## Authors’ Contributions

A Klein founded and leads the MindLogger project, led the present study, wrote the manuscript, and contacted all of the companies/organizations as part of the product comparison that led to Figures 1-3.

A Klein, A Krishnakumar, AB, and KK collected information for Figures 1-3 (detailed in Appendix 1). SSG and SAA lead the Repronim effort underlying MindLogger’s data schema.

A Krishnakumar helped to identify candidate features for MindLogger based on discussions with different stakeholders.

JC, BT, IS, NA, A Keshavan, and HR contributed to software engineering of MindLogger.

WVA helped to direct all aspects of MindLogger software development and design as MindLogger’s project manager.

KRM and JS contributed to the MindLogger project through content developed by the NIMH Family Study of Affective Spectrum Disorders.

YX helped to coordinate content in the NIMH-EMA, DBT, and KSADS applets, and found, modified, and designed images and icons for the NIMH-EMA, DBT, and EDUCATE applets.

SS conducted repeated, extensive quality assurance of MindLogger functionality.

AB and A Krishnakumar managed MATTER Lab volunteers in the US and in France.

LMA managed research collaborations as Research Operations Manager for the Child Mind Institute.

ABL and A Krishnakumar oversaw formulation of MindLogger requirements relevant to CRI projects. MM and SSG provided consultation.

KRM, JS, JC, LMA, WVA, BT, IS, NA, and MM reviewed and modified the manuscript for scientific content. AB helped format the manuscript.

## Disclaimer

The views and opinions expressed in this article are those of the authors and should not be construed to represent the views of any of the sponsoring organizations, agencies, or the US government.

## Multimedia Appendix 1

Supplementary materials.

Appendix1_product_search_protocol.pdf

## Abbreviations

API: Application Programmer Interface
DBT: Dialectical Behavior Therapy
EMA: Ecological Momentary Assessment
NIMH: National Institute of Mental Health

## References

1. Kessler RC, Berglund P, Demler O, Jin R, Merikangas KR, Walters EE. Lifetime prevalence and age-of-onset distributions of DSM-IV disorders in the National Comorbidity Survey Replication. Arch Gen Psychiatry 2005 Jun;62(6):593–602. PMID:15939837

2. Merikangas KR, He J-P, Brody D, Fisher PW, Bourdon K, Koretz DS. Prevalence and treatment of mental disorders among US children in the 2001–2004 NHANES. Pediatrics 2010 Jan;125(1):75–81. PMID:20008426

3. Demyttenaere K, Bruffaerts R, Posada-Villa J, et al. Prevalence, severity, and unmet need for treatment of mental disorders in the World Health Organization World Mental Health Surveys. JAMA 2004 Jun;291(21):2581–2590. PMID:15173149

4. Kohn R, Saxena S, Levav I, Saraceno B. The treatment gap in mental health care. Bull World Health Organ 2004 Dec;82:858–866. PMID:15640922

5. Saxena S, Thornicroft G, Knapp M, Whiteford H. Resources for mental health: Scarcity, inequity, and inefficiency. The Lancet 2007 Sep;370(9590):878–889. PMID:17804062

6. Raine R, Carter S, Sensky T, Black N. ‘Referral into a void’: Opinions of general practitioners and others on single point of access to mental health care. J R Soc Med 2005 Apr;98(4):153–157. PMID:15805555

7. Ward-Ciesielski EF, Rizvi SL. Finding mental health providers in the United States: A national survey and implications for policy and practice. J Ment Health 2019 Oct;1–7. PMID:31647364

8. Malowney M, Keltz S, Fischer D, Boyd JW. Availability of outpatient care from psychiatrists: A simulated-patient study in three U.S. cities. Psychiatr Serv 2014 Oct;66(1):94–96. PMID:25322445

9. E National Academies of Sciences, H and M Division, B on HC Services, and C to E the D of VAMH Services. Timely Access to Mental Health Care. US: National Academies Press; 2018.

10. Eisenberg D, Golberstein E, Gollust SE. Help-seeking and access to mental health care in a university student population. Med Care 2007 Jul;45(7):594–601. PMID:17571007

11. Lefebvre A, Sommerauer J, Cohen N, Waldron S, Perry I. Where did all the ‘no-shows’ go?*. Can J Psychiatry 1983 Aug;28(4). PMID:6627199

12. Campbell DGl, Downs A, Meyer WJ, McKittrick MM, Simard NM, O’Brien P. A preliminary survey of pediatricians’ experiences with and preferences for communication with mental health specialists. Fam Syst Health J Collab Fam Healthc. 2018 Sep;36(3):404–409. PMID:29199842

13. Sipe M. The Effects of Stigma toward Mental Illness on Family Physicians. Tucson, AZ: The University of Arizona; 2016.

14. Ginn SK Clark E. The medical profession and stigma against people who use drugs. Br J Psychiatry 2017 Dec;211(6):400–400. PMID:29196405

15. Klein JD, McNulty M, Flatau CN. Adolescents’ access to care: Teenagers’ self-reported use of services and perceived access to confidential care. Arch Pediatr Adolesc Med 1998 Jul;152(7):676–682. PMID:9667540

16. Kuramoto-Crawford SJ, Han B, McKeon RT. Self-reported reasons for not receiving mental health treatment in adults with serious suicidal thoughts. J Clin Psychiatry 2017 Jun;78(6):631–637. PMID:28406268

17. Gowen LK. Online mental health information seeking in young adults with mental health challenges. J Technol Hum Serv 2013 Apr;31(2):97–111. PMID:31742562

18. Dunster GP, Swendsen J, Merikangas KR. Real-time mobile monitoring of bipolar disorder: A review of evidence and future directions. Neuropsychopharmacology 2020 Sep. PMID:32919408

19. Saeb S, Zhang M, Karr SJ, et al. Mobile phone sensor correlates of depressive symptom severity in daily-life behavior: an exploratory study. J Med Internet Res 2015 Jul;17(7):e175. PMID:26180009

20. Trull TJ, Ebner-Priemer UW. Ambulatory assessment in psychopathology research: A review of recommended reporting guidelines and current practices. J Abnorm Psychol 2020 Jan;129(1):56–63. PMID:31868388

21. Granholm E, Loh C, Swendsen J. Feasibility and validity of computerized ecological momentary assessment in schizophrenia. Schizophrenia Bulletin 2008 May;34(3):507–514. PMID:17932087

22. Husky M, Gindre C, Mazure CM, et al. Computerized ambulatory monitoring in mood disorders: Feasibility, compliance, and reactivity. Psychiatry Research 2010 July;178(2):440–442. PMID:20488558

23. Johnson EI, Barrault M, Nadeau L, et al. Feasibility and validity of computerized ambulatory monitoring in drug-dependent women. Drug and Alcohol Dependence 2009 Jan;99(1-3):322–326. PMID:18692969

24. Johnson EI, Grondin O, Barrault M, et al. Computerized ambulatory monitoring in psychiatry: A multi-site collaborative study of acceptability, compliance, and reactivity. International Journal of Methods in Psychiatric Research 2009;18(1):48–57. PMID:19195050

25. Serre F, Fatseas M, Debrabant R, et al. Ecological momentary assessment in alcohol, tobacco, cannabis and opiate dependence: A comparison of feasibility and validity. Drug and Alcohol Dependence 2012 Nov;126(1-2):118–123. PMID:22647899

26. Husky M, Olié E, Guillaume S, et al. Feasibility and validity of ecological momentary assessment in the investigation of suicide risk. Psychiatry Research 2014 Dec;220(1-2):564–570. PMID:25155939

27. Moore RC, Swendsen J, Depp CA. Applications for self-administered mobile cognitive assessments in clinical research: A systematic review. International Journal of Methods in Psychiatric Research 2017 Dec:e1562. PMID:28370881

28. Cognition Measures. Health Measures. https://www.healthmeasures.net/explore-measurement-systems/nih-toolbox/intro-to-nih-toolbox/cognition. Accessed July 06, 2020.

29. Weintraub S, Dikmen SS, Heaton RK, et al. Cognition assessment using the NIH Toolbox. Neurology 2013 Mar;80(11):S54–S64. PMID:23479546

30. Technology. Neuroscape. https://neuroscape.ucsf.edu/technology/. Accessed July 07, 2020.

31. Cambridge Cognition. Cambridge Cognition. https://www.cambridgecognition.com. Accessed July 07, 2020.

32. Schueller SM, Begale M, Penedo FJ, et al. Purple: A modular system for developing and deploying behavioral intervention technologies. J Med Internet Res 2014 Jul;16(7):e181. PMID:25079298

33. Purple Robot. CBITs TECH. https://tech.cbits.northwestern.edu/purple-robot/. Accessed July 07, 2020.

34. Lattie EG, Schueller SM, Sargent E, et al. Uptake and usage of IntelliCare: A publicly available suite of mental health and well-being apps. Internet Interv 2016 May;4:152–158. PMID:27398319

35. IntelliCare. IntelliCare. https://intellicare.cbits.northwestern.edu. Accessed July 07, 2020.

36. Beiwe Research Platform. Beiwe Research Platform. https://www.beiwe.org/. Accessed July 07, 2020.

37. Torous J, Kiang MV, Lorme J, et al. New tools for new research in psychiatry: A scalable and customizable platform to empower data driven smartphone research. JMIR Mental Health 2016 May;3(2):e16. PMID:27150677

38. Bot BM, Suver C, Neto EC, et al. The mPower study, Parkinson disease mobile data collected using ResearchKit. Sci Data 2016 Mar;3(1). PMID:26938265

39. mPower 2.0. Parkinson mPower. https://parkinsonmpower.org/your-story. Accessed July 07, 2020.

40. ResearchKit and CareKit. Apple. http://www.apple.com/researchkit/. Accessed July 07, 2020.

41. ResearchStack. ResearchStack. http://researchstack.org/. Accessed July 07, 2020.

42. BIDMC - Division of Digital Psychiatry. Division of Digital Psychiatry Beth Israel Deaconess Medical Center. https://apps.digitalpsych.org/. Accessed July 07, 2020.

43. Torous JB, Chan SR, Yellowlees PM, et al. To use or not? Evaluating ASPECTS of smartphone apps and mobile technology for clinical care in psychiatry. J Clin Psychiatry 2016 Jun;77(6):e734–e738. PMID:27136691

44. App Evaluation Model. Psychiatry.org. https://www.psychiatry.org/psychiatrists/practice/mental-health-apps/app-evaluation-model. Accessed July 07, 2020.

45. MindLogger. Mindlogger. https://mindlogger.org/. Accessed July 06, 2020.

46. Lyon AR, Munson SA, Renn BN, et al. Use of human-centered design to improve implementation of evidence-based psychotherapies in low-resource communities: Protocol for studies applying a framework to assess usability. JMIR Res Protoc 2019 Oct;8(10):e14990. PMID:31599736

47. Vilardaga R, Rizo J, Zeng E, et al. User-centered design of learn to quit, a smoking cessation smartphone app for people with serious mental illness. JMIR Serious Games 2018 Jan;6(1):e2. PMID:29339346

48. Alexander LM, Escalera J, Ai L, et al. An open resource for transdiagnostic research in pediatric mental health and learning disorders. Sci Data 2017 Dec;4(1). PMID:29257126

49. Merikangas KR, Cui L, Heaton L, et al. Independence of familial transmission of mania and depression: Results of the NIMH family study of affective spectrum disorders. Mol Psychiatry 2014 Feb;19(2):214–219. PMID:24126930

50. Lamers F, Swendsen J, Cui L, et al. Mood reactivity and affective dynamics in mood and anxiety disorders. J Abnorm Psychol 2018 Oct;127(7):659–669. PMID:30335438

51. Merikangas KR, Swendsen D, Hickie IB, et al. Real time mobile monitoring of the dynamic associations among motor activity, energy, mood and sleep among adults with bipolar disorder. JAMA Psychiatry 2019 Feb;76(2):190–198. PMID:30540352

52. Johns JT, D.J, Merikangas K, et al. Fragmentation as a novel measure of stability in normalized trajectories of mood and attention measured by ecological momentary assessment. Psychol Assess 2019 Mar;31(3):329–339. PMID:30802118

53. Anguera JA, Jordan JT, Castaneda D, et al. Conducting a fully mobile and randomised clinical trial for depression: access, engagement and expense. BMJ Innov 2016 Jan;2(1):14–21. PMID:27019745

54. The Most Popular Database For Modern Apps. MongoDB. https://www.mongodb.com. Accessed July 06, 2020.

55. React Native . A Framework For Building Native Apps Using React. React Native. https://reactnative.dev/. Accessed July 06, 2020

56. The Progressive JavaScript Framework. VueJS. https://vuejs.org/. Accessed October 01, 2020.

57. Data-Driven Documents. D3.js. https://d3js.org/. Accessed October 01, 2020.

58. We Help You Stop Cyberattacks with Cybersecurity Consulting & Training. Alpine Security. https://alpinesecurity.com/. Accessed July 07, 2020.

59. Common Public Attribution License 1.0. Open Source Initiative. https://opensource.org/licenses/CPAL-1.0. Accessed July 07, 2020.

60. mindlogger-app. GitHub. https://github.com/ChildMindInstitute/mindlogger-app/blob/master/LICENSE.md. Accessed October 16, 2020.

61. Girder: A Data Management Platform — Girder 3.1.0 Documentation. Readthedocs. https://girder.readthedocs.io/en/stable/. Accessed July 06, 2020.

62. CherryPy — A Minimalist Python Web Framework. CherryPy. https://cherrypy.org/. Accessed October 01, 2020.

63. Amazon Web Services (AWS) - Cloud Computing Services. Amazon Web Services, Inc. https://aws.amazon.com/. Accessed July 06, 2020.

64. Cloud Computing Services. Google Cloud. https://cloud.google.com/. Accessed July 06, 2020.

65. Managed MongoDB Hosting | Database-as-a-Service. MongoDB. https://www.mongodb.com/cloud/atlas. Accessed July 06, 2020.

66. Redis. Redis. https://redis.io/. Accessed October 01, 2020.

67. Reproschema Documentation. Repronim. https://www.repronim.org/reproschema/. Accessed October 01, 2020.

68. for Linking Data. JSON-LD. https://json-ld.org/. Accessed October 01, 2020.

69. Moreau L, Missier P. PROV-DM: The PROV Data Model. W3C. https://www.w3.org/TR/2012/CR-prov-dm-20121211/diff.html. Accessed October 01, 2020.

70. Moats LC, Dakin KE. Basic facts about dyslexia and other reading problems. Baltimore: The International Dyslexia Association; 2008.

71. Duncan GJ, Magnuson K. Socioeconomic status and cognitive functioning: Moving from correlation to causation. Wiley Interdiscip Rev Cogn Sci 2012 May;3(3):377–386. PMID:26301469

72. Francis DA, Caruana N, Hudson JL, et al. The association between poor reading and internalising problems: A systematic review and meta-analysis. Clin Psychol Rev 2019 Feb;67:45–60. PMID:30528985

73. Becker N, Vasconcelos M, Oliveira V, et al. Genetic and environmental risk factors for developmental dyslexia in children: Systematic review of the last decade. Dev Neuropsychol 2017;42(7-8):423–445. PMID:29068706

74. Zeiss AM, Karlin BE. Integrating mental health and primary care services in the department of veterans affairs health care system. J Clin Psychol Med Settings 2008 Mar;15(1):73–78. PMID:19104957

