## Appendix 1 for "Remote Digital Psychiatry: MindLogger for Mobile Mental Health Assessment and Therapy"

### Appendix 1 for “Remote Digital Psychiatry: MindLogger for Mobile Mental Health Assessment and Therapy”: Protocol for product search supporting Figures 1-3

##### *Review criteria for existing customizable data collection tools*

In this Appendix, we outline the protocol we followed to gather information about products that appear to have desired primary features for conducting experience sampling studies. Specifically, the product must have an administrative user interface for creating and scheduling times and days for recurring, customized questionnaires, where end users receive and respond to scheduled notifications via an iOS or Android app on a mobile device. We cataloged other characteristics of interest for the products that satisfy the above criteria. For Google searches, we did not enclose any search queries with quotation marks unless explicitly indicated. We relied primarily on products’ official websites for information, and where they were unclear, we contacted the companies/organizations through their online contact forms or via email to clarify. For those products that satisfied our primary criteria, we engaged in product demonstrations and/or testing on our own devices.

**Search strategy**

1) Search for websites of candidate products identified by research collaborators and clinical colleagues over the course of the last three years.
2) Collect the first 20 Google search results (no ads) for each of three queries:
    a) “digital electronic data capture systems”
    b) “mobile phone software sensor data collection”
    c) “alternative to qualtrics”
3) Follow product leads in the search results. For example, if a website says “Top ten apps for...” then we conduct additional online searches for websites for each of the 10 apps that fit our inclusion criteria (below). We rely on official product websites, GitHub repositories of the creators of the products, or official apps on Apple’s App Store or Google Play. If the name of the project is too generic or has multiple words, we add the name of the company to the search or add quotation marks around the project name.
4) Include only those products that are currently in use.
    a) To be currently in use, product must have an active website, and any of the following:
        i) Website contains information for obtaining the product (“Download”, “Book a demo”, “Contact us to get a quote”, or a similar phrase)
        ii) Last release date was within the last two years (to October 10, 2020), and the most recent date from any of these sources:
            (1) Release notes (“release” or “version”) in product site’s news/blog
            (2) Release information in a GitHub open source repository
            (3) App Store “version info” or Google Play “last updated”
            (4) Google search: “[project name] last release”
            (5) Latest commit date in open source repository

5) Include only those products that have a mobile component (Android and/or iOS).
    a) Search website directly for “app” and “application”
    b) Search website using Google: “*site:<domain name> <keyword>*”,
    where [keyword] is any of “app”, “android”, “ios”, “store”, “google play store”, “itunes”, “app store”, “mobile”, “native”; *example: “site:[mindlogger.org] [app]”*
    c) Search website for links to Apple’s App Store or Google Play.
    d) Search Google: *“[product name] [keyword]”* using above keywords.
    (Apps in the app stores are not always mentioned in a product’s website.)

    Notes:
    1. “Mobile” specifically refers to “mobile app” or “for building mobile apps”; not, for example, “mobile data collection”.
    2. “Android application”, “Native mobile apps”, “Available on Apple (iOS) and Android devices”, and “supports ios and android devices” are all positive indicators.
    3. Phrases such as “works on desktop and mobile” and “compatible with android and ios devices” are not direct indicators, so follow any leads/links.
    4. Check in the product's list of software requirements.
    5. The end user product itself must be an app, and not for building apps, and search results need to clearly state that the product is a mobile/native app, not a web app that works on iOS/Android.

6) Include only those products that have an administrative user interface for creating and scheduling times for recurring, customized questionnaires, where end users receive and respond to scheduled notifications via an iOS and Android app.
    a) Look through the entire website for any information or links to information to confirm that there is a user interface (not just a code base or SDK) for creating customized questionnaires and for scheduling notifications to mobile app users to take these questionnaires.
    b) Where there is any ambiguity, contact the company/organization to clarify. Allow up to two weeks to receive a response from the company. If after an email exchange confirms that a product meets the primary criteria, schedule an online demonstration and request a free trial to explore the product. Include those products that meet the primary criteria.

7) Additional exclusion criteria -- we did not include products that:
    a) rely on SMS, email, or other modes of communication outside of the mobile app to send and receive notifications.
    b) were unwilling to demonstrate or provide a trial without a legal agreement.
    c) are intended for use by internal business employees of a company, not participants of a study or clinic, and require individual licenses or logins or fees per device.

**Additional information about each product**
To determine whether a product includes additional features of interest, we asked product developers or demonstrators some form of the following questions.

*Information presentation / data capture features*

1. When administrators use the product to create a customized questionnaire, can they include any of the following as means of presenting information to end users for a given question?:
    d) Image
    e) Audio clip
    f) Video clip
    g) Countdown/timer
    h) Conditional (branching/skip) logic
2. When administrators use the product to create a customized questionnaire, can they include any of the following to capture data from end users?:
    a. Text entry
    b. Camera photo
    c. Audio recording
    d. Video recording
    e. Drawing
    f. GPS location
3. Offline notifications: Does the product deliver local operating system notifications, where an end user receives notifications in their mobile device's notification bar at scheduled times? This is distinct from push notifications, which require an internet connection at the time that the notification is to be received, and from simple in-app notifications, which require the end user to be using the app to see their notifications.
4. When an end user taps on a notification in their mobile device's notification bar, does it take them directly to their scheduled activity within the app?
5. Does the product include access to additional sensors (e.g., accelerometer)?
6. When administrators use the product to create a customized questionnaire, how many languages can they choose from to use the administrative interface? This is distinct from how many different languages the end users can see.

*Privacy/security*

7. Does the product encrypt data on the mobile device, in transit, or on the server?
8. Is the product end-to-end encrypted, or would it be possible for the product creators to see respondents' response data?
9. Can an end user delete their own data without having to make a request from an administrator or the product creator?
10. Can an administrator delete an individual end user's data without having to make a request from the product creator?

*Access to the product, software, and data*

11. Can more than one end user access the app on the same mobile device, even if it means logging out and logging back in again?
12. Is there a data visualization dashboard to review individual end user's response data?
13. If there is a data visualization dashboard, can an administrator give someone access to review only one end users' response data in the dashboard?
14. Is there a consumer-facing application programming interface (API)?
15. Does the product include a software license?
16. Is the product open source?
17. Do you have to pay money to use the product for personal or research purposes?

**Preliminary products**

The 101 products below met the "Search strategy" criteria 1-5 above
(query matches for products that appear to be alive and mobile).
Products in bold are listed in the table below and in the main manuscript's Tables 1, 2, and 3:

| | |
|---|---|
| 123FormBuilder | https://www.123formbuilder.com/mobile-surveys/ |
| **Appbakery** | https://trialx.com/appbakery |
| AWARE | https://www.awareframework.com |
| B2W Inform | https://www.b2wsoftware.com/products/inform/ |
| **Beiwe** | https://www.beiwe.org/ |
| CAPI Survey Software | https://www.idsurvey.com/en/capi-software-face-to-face-interviews/ |
| Chesshealth | https://www.chess.health/solutions/ |
| Clinical Ink's Lumenis | https://www.clinicalink.com |
| Commcare | https://www.dimagi.com/commcare/ |
| Confirmit | https://www.confirmit.com/Products/Mobile/ |
| CRFweb app | https://crfweb.com/app/ |
| DADOS | https://www.dadosproject.com/solutions/#toggle-id-1 |
| DataCollector App | https://github.com/seemoo-lab/seemoo-mobile-sensing |
| DataScope | https://www.mydatascope.com/en |
| Delighted | https://delighted.com/ |
| Device Magic | https://www.devicemagic.com/ |
| **DFengage ePRO** | https://www.dfnetresearch.com/software/dfengage/ |
| Dharma | https://dharmaplatform.com/ |
| Dozuki | https://www.dozuki.com/solutions/need/work-instructions |
| Encapsia eSource app | https://encapsia.com/esource/ |
| Epicollect5 | https://five.epicollect.net/ |
| ePRO | https://www.crucialdatasolutions.com/epro-ecoa-econsent/ |
| **Ethica data** | https://ethicadata.com/ |
| **ExpiWell** | https://www.expiwell.com/ |
| FastField | https://www.fastfieldforms.com/ |
| FDA MyStudies | https://www.fda.gov/drugs/science-and-research-drugs/covid-mystudies-application-app |
| Flask | https://www.flaskdata.io/products/ |

| | |
|---|---|
| FME | https://www.safe.com/data-types/sensors/ |
| Folia | https://www.foliahealth.com/ |
| Form.com | https://www.form.com/platform/features/mobile-forms/ |
| FormAssembly | http://www.formassembly.com/ |
| FormFoundry | http://www.formfoundry.io/ |
| Forms on Fire | https://www.formsonfire.com/features/ |
| Formstack | https://www.formstack.com/online-forms |
| Fulcrum | https://www.fulcrumapp.com/ |
| Gather | https://gathercapture.com/data-capture/custom-forms |
| GoCanvas | https://www.gocanvas.com/ |
| GoFormz | https://www.goformz.com |
| GoSpotCheck | https://www.gospotcheck.com/ |
| GoSurvey | https://www.gosurvey.in/ |
| **Ilumivu's mEMA** | https://ilumivu.com/solutions/ecological-momentary-assessment-app/ |
| INKWRX | https://www.inkwrx.com/ |
| Inquisit | https://www.millisecond.com/products/inquisit6/inquisitmobile.aspx |
| iQapture | https://valuechain.com/iqapture#features |
| Klipboard | https://klipboard.io/feature/workflow-generator-pdf-designer/ |
| KoboToolbox | https://www.kobotoolbox.org/ |
| Kordata | https://www.kordata.com |
| LabFront | https://www.physioq.org/about |
| **LifeData's RealLife Exp** | https://www.lifedatacorp.com/solutions-research/ |
| M-sense | https://www.m-sense.de/en/ |
| Magpi | https://www.magpi.com/uses |
| MathWorks Sensor Data Collection | https://www.mathworks.com/help/matlabmobile_android/sensor-data-collection.html |
| Medallia Ask Now | https://www.medallia.com/platform/ask-now/ |
| Medic Mobile | https://medicmobile.org/ |
| **Metricwire** | https://metricwire.com/ |
| MindLAMP 2 | https://www.digitalpsych.org/lamp.html |
| MobileCoach | https://www.mobile-coach.eu/ |
| Mopinion | https://www.mopinion.com |
| movisensXS | https://xs.movisens.com |
| naturalForms | https://www.web.naturalforms.com/mobile-form-builder-app |
| Nebu Dub InterViewer | https://www.nebu.com/nebu-dub-interviewer-questionnaire-design-and-survey-programming |
| Nintex App Studio | https://www.nintex.com/workflow-automation/mobile-apps/ |
| ODK (Open Data Kit) | https://getodk.org/, https://docs.getodk.org/collect-intro/ |
| **Open HealthHub Improve** | https://www.openhealthhub.com/en/products/improve-mobile-app/ |
| Overlap | https://www.overlaphealth.com/ |
| PIEL Survey app | https://pielsurvey.org/ |
| Pisano | https://www.pisano.co/en |
| Poimapper Plus app | https://www.poimapper.com/en/features/ |
| Poll Everywhere | https://www.polleverywhere.com |
| Pollscape | https://pollscape-d2b4b.firebaseapp.com/ |
| Pronto forms | https://www.prontoforms.com/ |
| PsychSurveys | http://psych-surveys.com/ |
| PSYT | https://www.psyt.co.uk/ |
| Qualaroo | https://qualaroo.com/ |
| QuestionPro | https://www.questionpro.com/ |

| | |
|---|---|
| QuickTapSurvey | https://www.quicktapsurvey.com/features |
| RADAR-Base aRMT app | https://radar-base.org/index.php/getting-started-with-radar-base/try-out-armt-app/ |
| REDCap MyCap | https://projectmycap.org/ |
| Repsly | https://www.repsly.com |
| Science 37 | https://www.science37.com/ |
| Secutrial | https://www.secutrial.com/en/advantages/ |
| SensingKit | https://www.sensingkit.org/ |
| Sensors Data Collector | https://play.google.com/store/apps/ details?id=com.opendata.labs.sensorsdatacollector&hl=en |
| Sensus | https://predictive-technology-laboratory.github.io/sensus/ |
| Snap Mobile Anywhere | https://www.snapsurveys.com/survey-software/snap-mobile-anywhere/ |
| SoGoSurvey | https://www.sogosurvey.com/mobile-app/ |
| Spokk | https://www.spokk.io/ |
| SurveyCTO | https://www.surveycto.com |
| SurveyLab | https://www.surveylab.com/ |
| SurveyMonkey | https://www.surveymonkey.com/mp/mobile-surveys/ |
| SurveyPocket | https://www.surveyanalytics.com/mobile |
| SurveySparrow | https://surveysparrow.com/survey-tools/online-survey-tools/ |
| Teamscope | https://www.teamscopeapp.com/ |
| Trayt | https://trayt.io/ |
| Trial Online ePro app | https://www.trialonline.se/epro-subject-app/ |
| UCLA MDS (ohmageX) | https://oit.ucla.edu/mobile-web-strategy/mobile-data-collection |
| unforgettable.me | https://www.unforgettable.me/ |
| Voxco Mobile | https://www.voxco.com/ |
| Xolomon | https://www.xolomon.com/research-institutes-foundations/?lang=en |
| Zenput | https://www.zenput.com/platform |
| **Zest** | https://zestmeup.com |
